# NGPINT: A Next-generation protein-protein interaction software

**DOI:** 10.1101/2020.09.11.277483

**Authors:** Sagnik Banerjee, Valeria Velásquez-Zapata, Gregory Fuerst, J. Mitch Elmore, Roger P. Wise

## Abstract

Mapping protein-protein interactions at a proteome scale is critical to understanding how cellular signaling networks respond to stimuli. Since eukaryotic genomes encode thousands of proteins, testing their interactions one-by-one is a challenging prospect. High-throughput yeast-two hybrid (Y2H) assays that employ next-generation sequencing to interrogate cDNA libraries represent an alternative approach that optimizes scale, cost, and effort. We present NGPINT, a robust and scalable software to identify all putative interactors of a protein using Y2H in batch culture. NGPINT combines diverse tools to align sequence reads to target genomes, reconstruct prey fragments and compute gene enrichment under reporter selection. Central to this pipeline is the identification of fusion reads containing sequences derived from both the Y2H expression plasmid and the cDNA of interest. To reduce false positives, these fusion reads are evaluated as to whether the cDNA fragment forms an *in-frame* translational fusion with the Y2H transcription factor. NGPINT successfully recognized 95% of interactions in simulated test runs. As proof of concept, NGPINT was tested using published data sets and recognized all validated interactions. NGPINT can be used in any organism with an available reference, thus facilitating the discovery of protein-protein interactions in non-model organisms.

## INTRODUCTION

All organisms respond to stimuli through a network of interacting proteins and other biomolecules. Such interactions play a pivotal role in biological processes such as signal transduction (Chen et al., 2016), gene transcription (Blaskovich et al., 2003), protein translation (Gumbart et al., 2009), disease regulation (Wang et al., 2012; Sharma et al., 2015) and developmental control (Braun and Gingras, 2012). It has been hypothesized that more than 80% of proteins form complexes with other proteins to carry out their respective functions (Berggård et al., 2007). Indeed, investigations into protein-protein interactions (PPI) are critical to evaluate information flow within the cell (Vinayagam et al., 2011; Nietzsche et al., 2016). Accordingly, PPI datasets (Arabidopsis_Interactome_Mapping_Consortium, 2011; Yu et al., 2011; Rolland et al., 2014) have been used extensively to construct interaction databases to promote the understanding of protein function and regulation (Hermjakob et al., 2004; Oughtred et al., 2019; Szklarczyk et al., 2019).

A multitude of *in vitro* and *in vivo* biochemical techniques have been designed to investigate and analyze binary PPI (Rao et al., 2014; Zhou et al., 2016). Commonly used methods include co-immunoprecipitation (Phizicky and Fields, 1995), protein pull down (Kaelin Jr et al., 1991), fluorescence and bioluminescence resonance energy transfer (Zal and Gascoigne, 2004; Rainey and Patterson, 2019), protein-fragment complementation assay (Galarneau et al., 2002; Remy and Michnick, 2006; Morell et al., 2009), and yeast two-hybrid (Walhout and Vidal, 2001; Vidal and Fields, 2014). These PPI assays are binary, *i.e*., they are used to interrogate a single pair of proteins, and thus demand significant time and resource commitment (Venkatesan et al., 2009).

Among PPI detection methods, yeast two-hybrid (Y2H) has been widely used (Walhout and Vidal, 2001; Vidal and Fields, 2014). In Y2H, each protein of interest is expressed as a translational fusion to different domains of a transcription factor (TF), typically GAL4 or LexA. One protein (the “bait”) is fused to the DNA binding domain (DBD) and the second protein (the “prey”) is fused to the transcriptional activation domain (AD). If these two fusion proteins interact in yeast, reconstitution of transcription factor activity (DBD+AD) results in expression of reporter genes. Common reporters encode proteins that are required for yeast growth on selective media or used for colorimetric assays. To discover novel interactions with no prior knowledge, Y2H can be adapted to screen a single bait against a prey cDNA library. Traditionally, this process has been limited to picking individual yeast colonies that grow on selective media and using Sanger sequencing to identify the putative interacting partners. Careful follow-up experiments must be performed to identify false positives, *i.e*., preys that can auto-activate the reporters in the absence of a *bona fide* PPI with the bait protein.

The falling costs of next-generation sequencing have enabled more recent high-throughput approaches to PPI discovery, collectively termed next-generation interaction screening (NGIS) (Suter et al., 2015). These methods employ deep sequencing to score the output from Y2H screens and can be scaled to process multiple interactions in parallel (Lewis et al., 2012; Weimann et al., 2013; Pashkova et al., 2016; Trigg et al., 2017; Erffelinck et al., 2018; Kessens et al., 2018; Yang et al., 2018; Zong et al., 2020). Generally, NGIS involves performing Y2H selection in batch and pooling positive yeast colonies after reporter selection. Prey sequences are amplified by PCR and the resulting amplicons are analyzed by next-generation sequencing. The resulting reads enable identification of the prey cDNA sequences and read counts are used to quantify abundance of each cDNA relative to a control bait and/or non-selected growth conditions in the yeast populations.

Model organisms such as human (Yu et al., 2011) and Arabidopsis (Trigg et al., 2017) have the advantage of full-length ORF libraries to facilitate genome-scale NGIS. However, most organisms outside models do not have available ORF resources and ORF libraries do not necessarily reflect the transcriptional landscape of specific conditions under study. Hence, the use of custom cDNA libraries derived from specific organisms and/or particular conditions of interest can be advantageous for large-scale PPI discovery in model and non-model organisms (Kessens et al., 2018; Zong et al., 2020).

In this report, we present NGPINT, a fully automated software platform to select candidate PPI from high-throughput Y2H-NGIS experiments. Unlike previously described pipelines (Pashkova et al., 2016), NGPINT can process data from any organism with an available genome and/or a transcriptome reference. The software accepts a single csv file from the user containing the location of the fastq read files, output directory, Y2H plasmid sequences, and the genome and/or the transcriptome to use for read mapping. The pipeline aligns the reads, recognizes fusion reads from the alignments, trims Y2H plasmid sequence from fusion reads, generates gene counts, computes statistics on differential abundance, and automates the identification of putative interacting partners that are expressed as *in-frame* translational fusions. Finally, a report is generated which compiles the results of all analyses into a single file. The output from the pipeline facilitates the selection of high confidence genes/transcripts for further downstream validations.

Using simulation and an experimental dataset, we assessed the success of NGPINT. NGPINT was able to deliver consistent performance recognizing over 95% of the simulated interactions with minimal false positives. Using recently published data (Erffelinck et al., 2018), use of NGPINT revealed all previously validated interactions. We also show that errors in base calls do not influence the detection of PPI. The software package can be accessed from https://github.com/Wiselab2/NGPINT released under MIT license and should be valuable to researchers performing NGIS to discover PPI in model and non-model organisms.

## IMPLEMENTATION of NGPINT

### Overview

The NGPINT architecture comprises several features including trimming reads, aligning reads to a genome and/or transcriptome, read counting, detection of fusion reads, and calculation of differential abundance. To facilitate smooth operation among all software that NGPINT relies on, users are recommended to set up a conda environment that offers a virtual local environment and does not interfere with other conflicting software packages. The software has been developed in a modular style to allow future upgrades. The core of the software has been coded in Python; this enables it to be deployed cross-platform, from a personal computer to a cloud-based server. Should the situation arise, the pipeline is designed to restart runs from the previous point of failure, eliminating the need to execute from the beginning. Additionally, most of the software modules have been parallelized to improve resource utilization and execution times.

A flow chart outlining the data processing and analysis steps are presented in (**Figure 1**). In its current form, the workflow is optimized for use with baits mated individually to a prey cDNA library under selected and non-selected growth conditions. Processing begins with removal of adapter sequences using Trimmomatic (Bolger and Giorgi, 2014). Trimmed reads are aligned to the reference with an initial run of STAR (Dobin et al., 2013) that aids the detection of fusion reads. A custom python script is used to trim Y2H plasmid sequence from the fusion reads, and then a second round of STAR is used to map the trimmed reads to the reference genome/transcriptome. Gene counts are obtained using Salmon (Patro et al., 2017) and DESeq2 (Love et al., 2014) is used for differential abundance analysis. Fusion read alignments are restructured and tagged which allows them to be differentiated from other kinds of reads when viewed in genome browsers. Finally, a file is generated that can assist users to construct primers for secondary validation of candidate PPI. Even though most of the modules are available individually, the implementation of those modules is tailored to address the unique aspects of Y2H data. NGPINT is fully automated to serve bench researchers with limited experience in the analysis of large-scale data sets.

**Figure 1.**
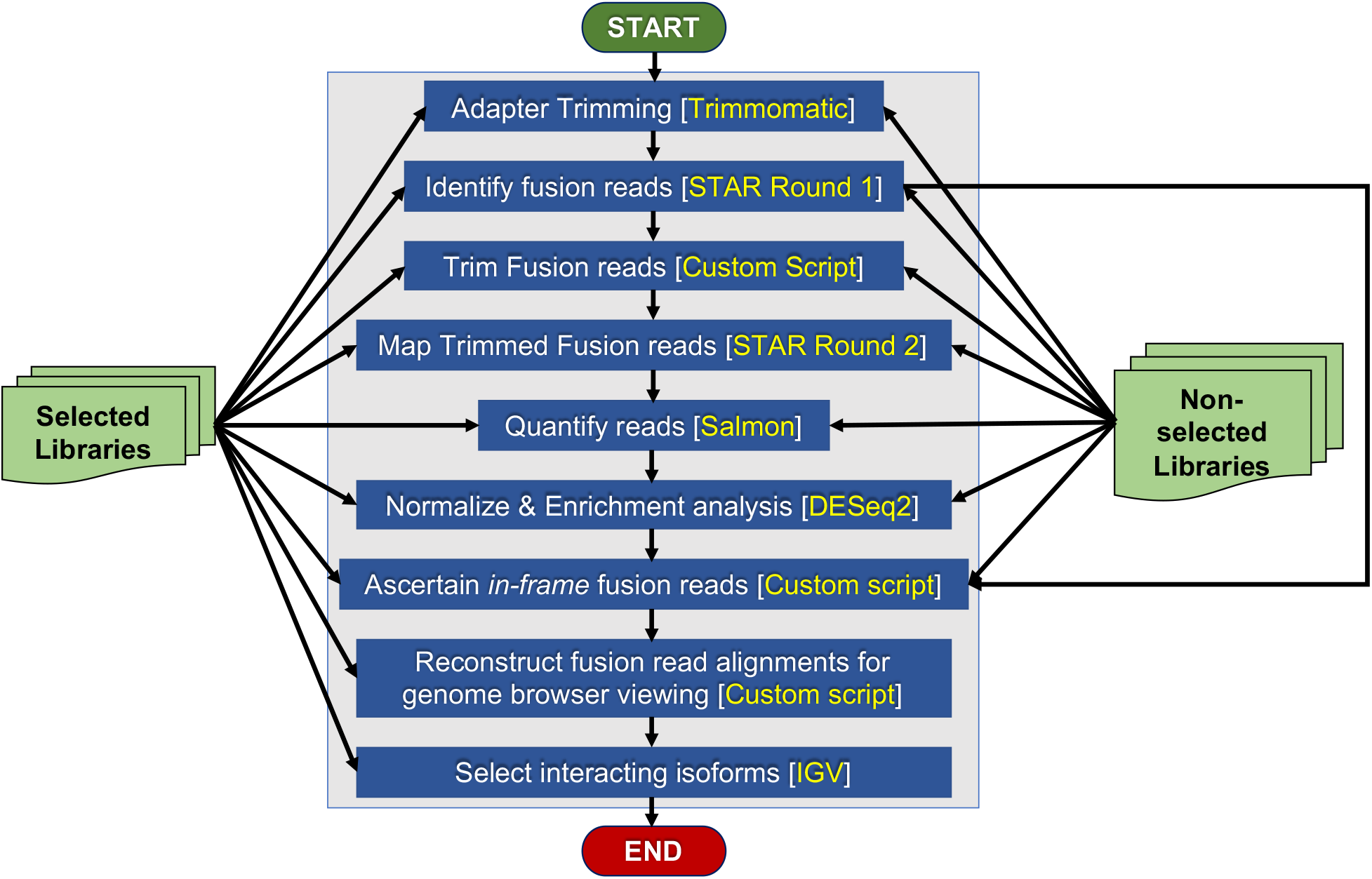
Workflow of NGPINT. Interaction between the modules of NGPINT has been illustrated in this workflow diagram. Raw reads from selected and non-selected samples are processed an automated fashion to output potential interactors. Processes are depicted using blue colored boxes and the green colored boxes represents the input data. The arrows connecting two processes indicate flow of both data and order of execution. Arrows from data boxes (those in green color) illustrate flow of information.

Central to this pipeline is the detection of Y2H plasmid/organism cDNA fusion reads that provide crucial information about the reading frame of the fusion protein expressed in yeast. Due to the stochastic nature of cDNA library preparation, detection of *in-frame* expression is essential to ensure that the native peptide sequence has been expressed as a translational fusion to the activation domain. A detailed description of each step is provided below:

### I. Inputs and Setup

All inputs to the pipeline are accepted in the form of a comma separated value (csv) configuration file (**Supplementary Table 1**). A sample csv file is provided with the software application package. The genome is expected to be a single fasta file and the associated transcriptome annotations should be provided in GTF format. If a genome is unavailable, a transcriptome fasta file can be provided in its place. In addition, the pipeline requires the Y2H plasmid sequences of both the ‘prey’ and the ‘bait’ plasmids. Oligonucleotide primer sequences used for amplification of preys are supplied as anchors to detect fusion reads. NGPINT requires indexing for mapping short reads to the reference and the Y2H plasmid sequences. Users have the option of generating the genome index for read mapping prior to execution and this step only needs to be completed once. Index generation speeds up the execution considerably, especially for large genomes (> 3 Gb). Users are encouraged to use the pipeline to generate the indices for at least one bait and then reuse the same index for analyzing additional baits. Finally, fastq files, for each replicate, corresponding to the selected and the nonselected conditions need to be provided.

#### A. Validating inputs

The pipeline starts by verifying all inputs. It checks for the presence of input files and ensures that all the files are in the appropriate formats. Checks are also performed to ensure that the requested CPUs and RAM are available. Genome indices used by the STAR aligner are generated if they are not provided by the user. Errors generated in this process are reported in the progress.log file in the output directory.

#### B. Adapter trimming

Adapter sequences are removed using Trimmomatic (Bolger and Giorgi, 2014). All adapter sequences required for trimming are provided with the software release. Custom adapters can be added if desired. No quality trimming is performed because of the improvement in sequencing technology rarely produces any incorrect bases (Pfeiffer et al., 2018). Also, most aligners allow for soft clipping of poor quality unmappable bases, thus eliminating the need to trim low-quality bases. Illumina sequencers sometimes produce reads which have trailing and/or leading ‘N’. Reads that are flanked with ‘N’ introduce ambiguity, which are soft clipped by STAR. During *in-frame* detection, soft-clipped portions of reads are compared with plasmid sequence. The presence of ‘N’s will increase hamming distance of the trimmed portion preventing it to be detected as a fusion read. Hence, such sequences are trimmed due to possible interference with detection of *in-frame* fusion reads.

#### C. Mapping to reference

Adapter-trimmed reads are aligned to the reference genome and to the Y2H vector sequences using two cycles of STAR (Dobin et al., 2013). STAR version 2.7.3a is included in the software package. In the first round, STAR is configured to allow at least 30% of a read to map. The low threshold of 30% ensures the capture of as many fusion-reads as possible. STAR can map reads to the genome and simultaneously transfer the alignments to the transcriptome (if provided), thus speeding up the operation of the entire pipeline. Reads that align to the genome and to the flanking vector sequence with soft clips are mined for fusion reads. In the second round of alignment with STAR, only the identified fusion reads that aligned to the flanking vector sequence in the first round are re-mapped after trimming the flanking sequence. Lastly, alignments from the two rounds of STAR are merged together.

### II. Detection of Fusion Reads

After performing batch Y2H and plasmid extraction, vector primers on either side of the cDNA insert are used to amplify prey sequences in the selected and non-selected yeast populations. (**Supplementary Figure S1A**). Sequence reads derived from both the flanking region of the Y2H plasmid and the prey cDNA are referred to as fusion reads (**Supplementary Figure S1B**). Reads that map to the reference but do not possess flanking plasmid sequence are referred to as fusion-free reads (**Supplementary Figure S1C**). The junction sequence at the 5’ end of the amplicon provides information essential to detect *in-frame* fusion.

Before mapping fusion reads to the reference, NGPINT **Algorithm 1** is first used to trim away the plasmid sequence, and the remaining part of the read is flagged. Reads that contain only plasmid sequence are discarded because they should not map to transcripts derived from the experimental organism(s). Fusion reads containing 3’ vector and poly-A tails, longer than half the length of the read, are also discarded since they do not contain useful information for mapping. Fusion reads are detected from alignments that contain soft clips. If the majority of a fusion read is made of cDNA, then it will align to the organism reference and the flanking region will be soft clipped. The remaining fusion reads that contain longer flanking region will map to the plasmid sequence and the region corresponding to the organism reference will be soft-clipped. Aligners typically generate a string (termed CIGAR) for each alignment to indicate matches, mismatches, insertion and deletion of nucleotides between the read and the reference segment (https://genome.sph.umich.edu/wiki/SAM). The cigar string of read alignment, that bear soft-clips, is used to determine the exact junction between the vector and the cDNA in fusion reads. Reads are then split at this junction, and the trimmed reads are mapped back to the genome. These reads will be later used to detect *in-frame* fusions between the TF activation domain and the cDNA. If the 5’ primer for prey amplification is designed too far away from the junction, some reads may have a very small portion of the cDNA sequence post-trimming. Such a small trimmed sequence (<25 bp) usually has multiple hits to the organism reference. To address this ambiguity, paired-end libraries can be used to facilitate mapping the mate-pair to the reference. NGPINT restructures the alignments which enables examination of the fusion reads along with all other reads in a genome browser such as Integrated Genome Viewer (IGV), with the flanking plasmid sequence soft-clipped (**Figure 2**). Visualization in this manner is useful for confirming the exact cDNA fragments that were expressed from the Y2H expression plasmid.

**Figure 2.**
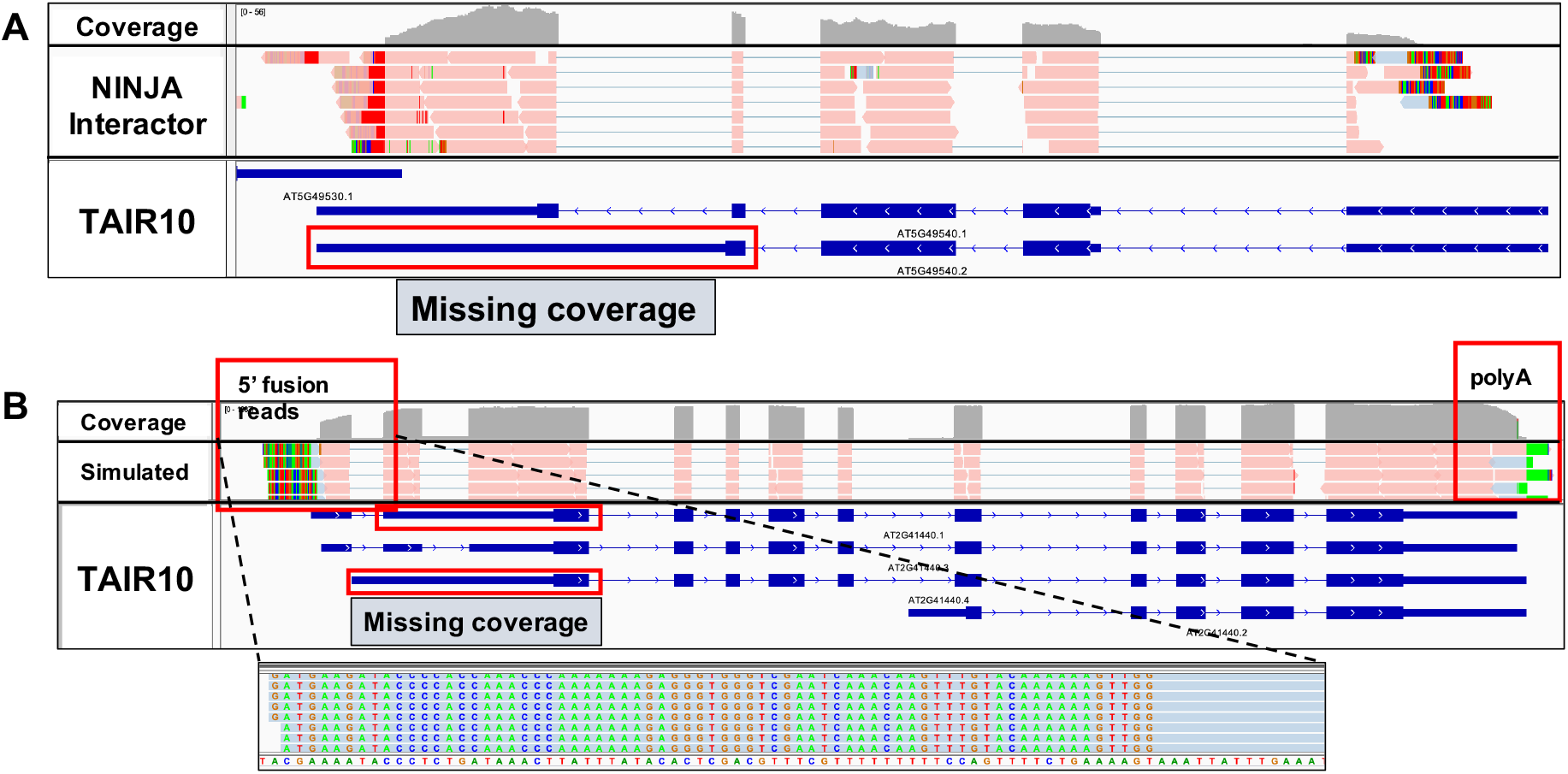
Selective protein isoforms interact with the bait. Integrated genome viewer (IGV) was used to visualize read coverage of genes and their isoforms. NGPINT generates alignment files which can be loaded into IGV to select the isoforms that have missing coverage. (**A**) Out of two isoforms only one protein isoform is detected as enriched with the Arabidopsis thaliana protein NINJA (AT4G28910). Absense of read coverage on the last exon of transcript isoform *AT5G49540.2* clearly indicate that it was not detected (**B**) A single isoform was selected to be overexpressed and interacting with the simulated bait. 5’ fusion reads and polyA tail flanks the simulated overexpressed transcript. Fusion reads are shown in red and fusion-free reads are blue. Magnifying the 5’ region will enable viewing the junction region in nucleotide level precision.

**Algorithm 1.**
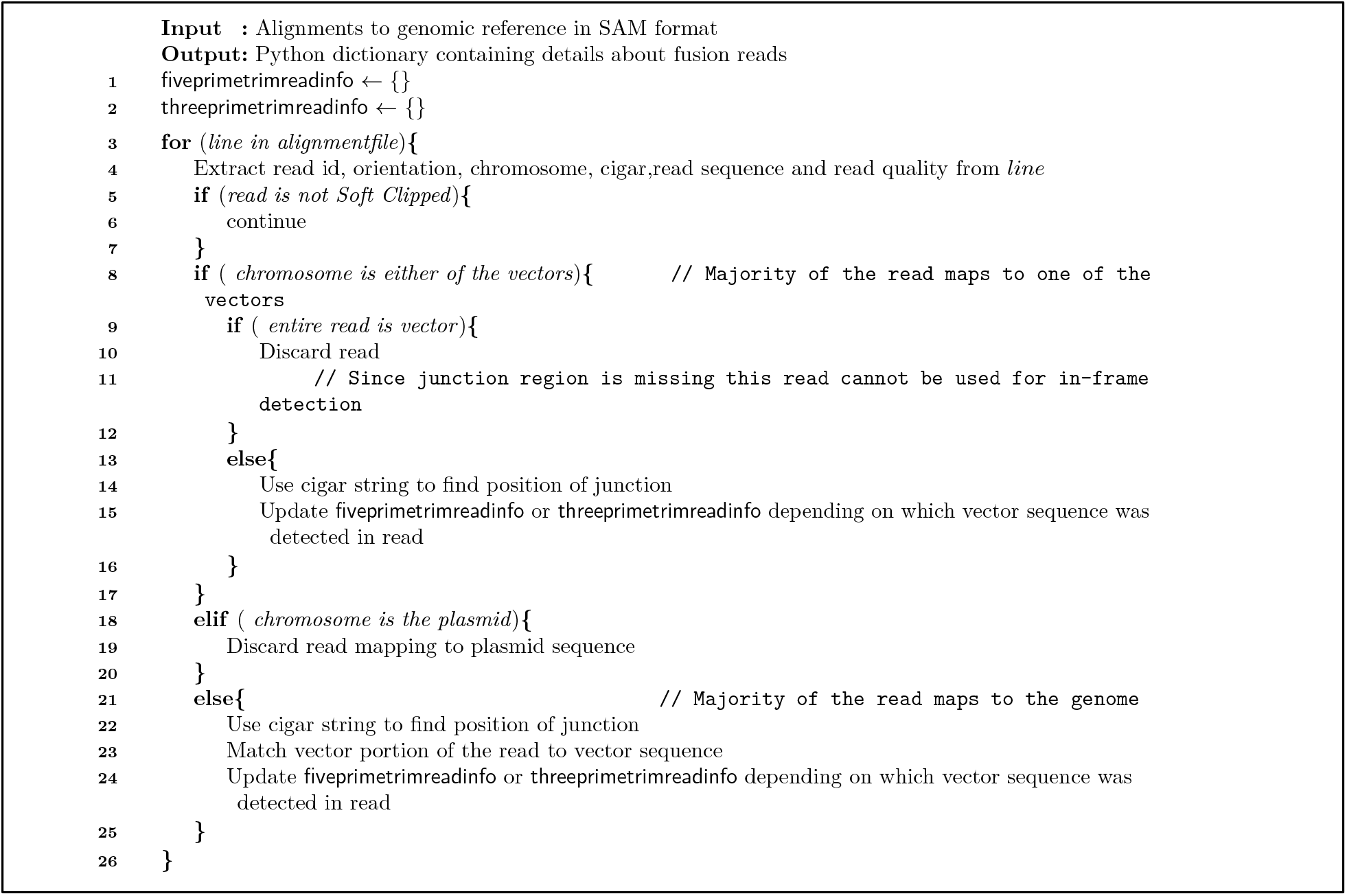
Step-by-step protocol to trim vector portion from fusion reads

#### A. Estimating read counts

Read counts are estimated using Salmon (Patro et al., 2017) in alignment-based mode. Trimmed fusion reads and fusion-free reads aligned to the reference sequences are provided to Salmon as input. In addition to the alignment files, a transcript to gene map is also provided when genome and transcriptome are used as a reference. Salmon employs statistical models to account for sample-specific parameter effects and guard against GC bias. Salmon is executed on multiple cores to speed up the counting process. NGPINT will generate counts for a single replicate, or multiple replicates, depending on the user’s experimental design, for both the selected and non-selected conditions.

#### B. Computing differential abundance

The NGPINT pipeline calculates differential abundance of each gene between the selected and the non-selected conditions using DESeq2 (Love et al., 2014). DESeq2 software was selected for this step because the Y2H-NGIS read counts resemble a negative binomial distribution (Pashkova et al., 2016). This analysis provides a relative measure to the enrichment levels of cDNAs after Y2H reporter selection, thus indicating a candidate PPI. The default median-of-ratios normalization (Anders and Huber, 2010) that is implemented in DESeq2 is not suitable for Y2H-NGIS because it assumes that only a small portion of the identified genes are differentially enriched (Velásquez-Zapata et al., 2020). Thus, raw read counts, ascertained by Salmon, were normalized using size factors estimated by library size (Dillies et al., 2013) prior to DESeq2 analysis.

#### C. Detecting in-frame fusion reads

In addition to identifying differentially enriched genes, it is advantageous to ensure that the gene expression occurred *in-frame* with the TF. Reverse transcription using poly-A tail capture ensures priming of most expressed transcripts even if they are partial. Partial cDNA fragments and full-length cDNA containing 5’ UTR are not guaranteed to be *in-frame* with the coding region of the TF-AD.

A potential solution is to construct cDNA libraries in all three reading frames. This ensures that all fragments (even the ones with 5’ UTR sequence) have the potential to be expressed as the native peptide sequence in one reading frame. Three-frame libraries can reduce false negatives but can also increase false positives by creating spurious peptide sequences, which might coincidentally interact with the ‘bait’ protein or be enriched for unidentified reasons. Hence, whether one selects a single- or three-frame library, detecting the reading frame of enriched cDNAs is critical.

Fusion reads mapped to each transcript are used to assess whether the transcript was expressed in the correct reading frame using NGPINT **Algorithm 2**. Transcript sequences, along with their CDS information, are extracted using the gffread utility of cufflinks (Trapnell et al., 2012). The cigar string of the alignment of each fusion read is interrogated to find the bases that are soft clipped and the location of mapping within the transcript. The CDS start location, the mapping location and the number of soft clipped bases are used to determine if the transcript was expressed *in-frame*. If the distance between the mapped location and start of the CDS is a multiple of three, the read is declared to be *in-frame* with the CDS. Then, the fraction of fusion reads aligning to transcripts *in-frame* with the coding sequence can be used to prioritize interactions for secondary validations (Velásquez-Zapata et al., 2020). Samtools (Li et al., 2009) was used to perform operations on alignment files and is included with NGPINT. Alignments of fusion reads to the genome can be viewed in any genome browser (**Figure 2B**).

**Algorithm 2.**
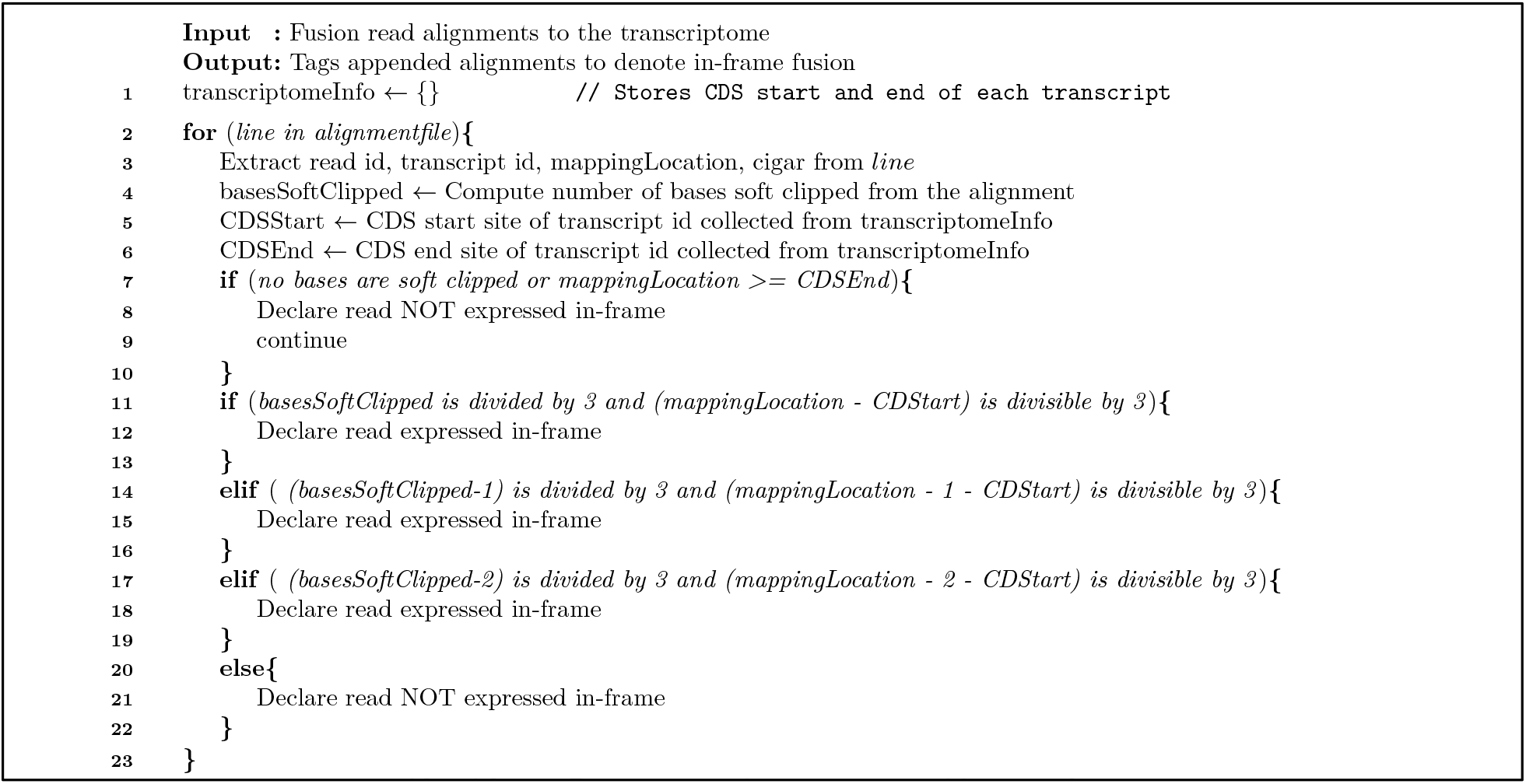
Determining in-frame expression of fusion reads

#### D. Generating visualizations for isoform selection and protein domain analysis

Once cDNAs that are differentially enriched and expressed *in-frame* have been identified, secondary validation should be conducted to confirm interactions. Protein-coding genes in eukaryotes often have multiple transcript isoforms, which encode different peptide sequences. These protein isoforms can have significant variability in their amino acid content and contain alternate domain structure, and thus, it is possible that only a few of the isoforms interact with the target protein. An example is illustrated in (**Figure 2A**), where it is clear that out of two possible transcript sequences only one was captured during cDNA synthesis. To facilitate visualization of the junction between vector and cDNA, NGPINT **Algorithm 3** is applied to modify the mappings of fusion reads to accommodate the previously trimmed vector sequence. Details about viewing the soft-clipped alignments can be found in the manual. When multiple cDNA fragments are identified from a single gene, these visualizations are useful to narrow down the minimal protein-coding region of the interacting prey (**Figure 3**).

**Figure 3.**
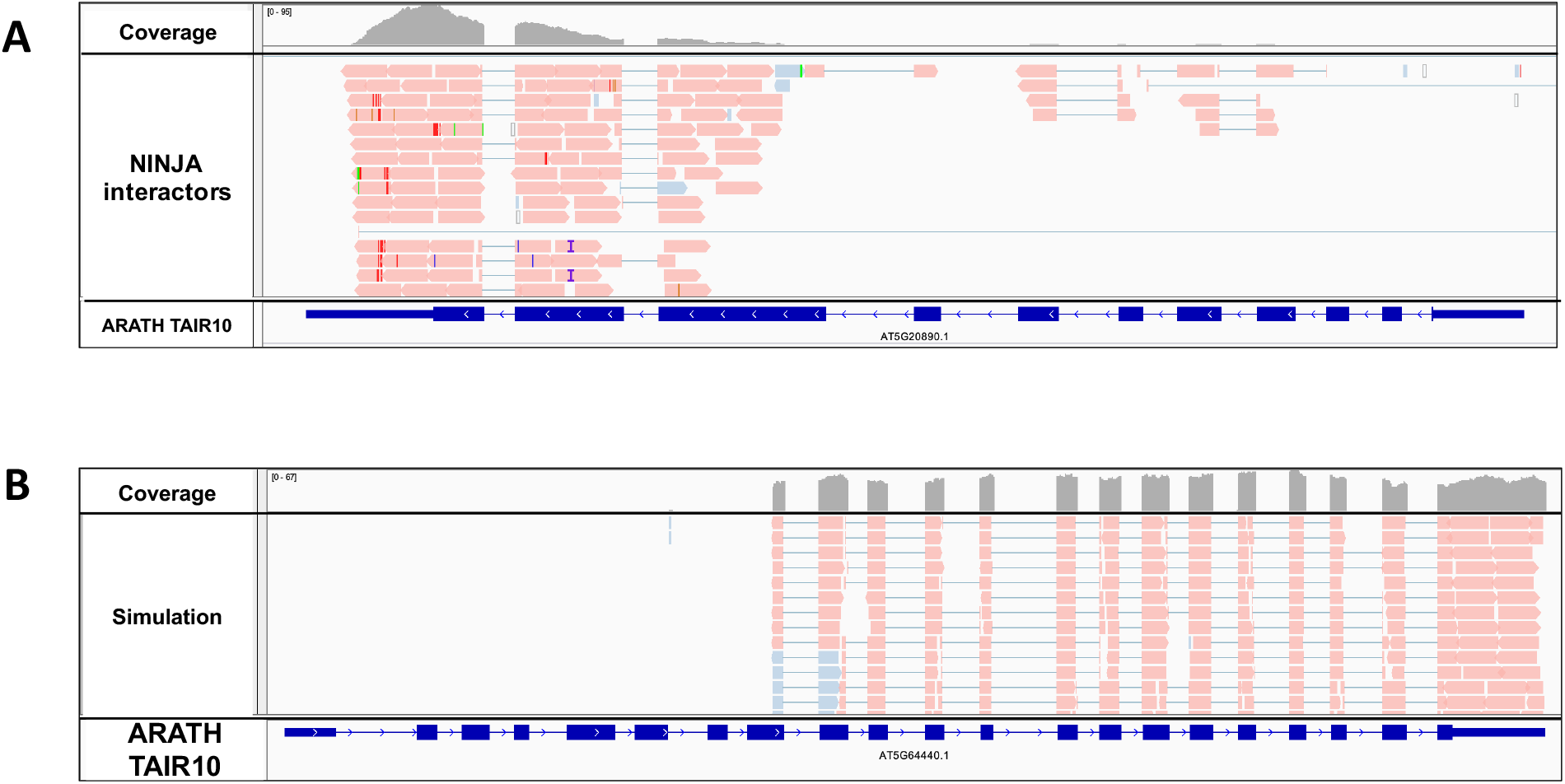
Domain specific interaction. (**A**) NINJA interacts with a portion of an Arabidopsis protein. Such cases help locate the domain of interaction and allow for a much more targeted study. (**B**) Similar cases were also simulated and was recognized by NGPINT.

**Algorithm 3.**
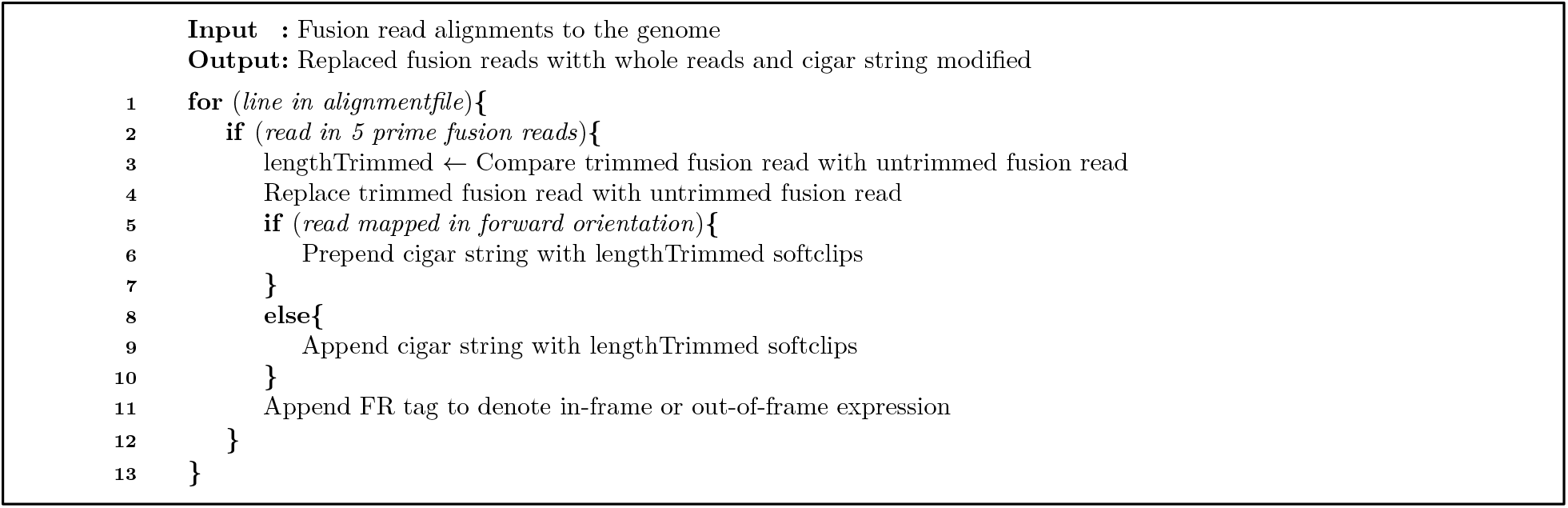
Preparing file for viewing in genome browser

### III. Pipeline Outputs

NGPINT generates a report file with all details about the mapping process, number of fusion reads recognized in each sample, and the duration of execution of each step. It also compiles a file with details about each transcript outlining the number of reads recognized *in-frame* for that transcript, statistics on differential abundance including fold change, p-values, and false discovery rate (FDR), location of any stop codons in the UTRs and transcripts per million (TPM) generated by Salmon (Patro et al., 2017). TPM uses the gene size to normalize counts accounting for the fact that longer genes are expected to produce more reads. We also have developed a companion module to prioritize candidate PPI using the Y2H-SCORES statistical workflow (Velásquez-Zapata et al., 2020). A wrapper has been included such that this statistical framework can be used directly from the NGPINT output and metadata files. Y2H-SCORES comprises a ranking system for the preys with three elements: 1) an enrichment score calculated from the selection/non-selection contrast, 2) when multiple baits have been screened, a specificity score defined as the degree of enrichment in pairwise comparisons of all baits under selection, and 3) an *in-frame* score calculated from the *in-frame* prey selection with the AD of the upstream transcription factor. Velásquez-Zapata and associates (2020) demonstrate that this scoring system efficiently ranks high-confidence interactors and is superior to differential abundance analysis alone.

Finally, a file is generated which contains sequence information useful for follow-up experiments. Users are recommended to use this file to design primers for cloning genes and gene fragments for further confirmation via binary assays. While it is important to view reconstructed fusions along with the transcript, most genome viewers do not offer the functionality of copying the CDS portion of the genome sequence. Hence, NGPINT generates an additional text file to facilitate primer design (**Figure 4**). Each transcript is represented by a header, its nucleotide sequence and a sequence of symbols. The header contains the gene identifier, sample origin, and various statistics from the differential abundance analysis. Coding sequences are represented in block letters and UTR in small letters. Three symbols are used {‘*’,’X’,’ ’} to depict whether a fusion read originated from that loci. If no fusion reads map to a nucleotide then it is represented as a space. Nucleotides from where *in-frame* fusion reads originate, have been annotated with a ‘*’. An ‘X’ has been put beneath nucleotides in the 5’ UTR from which fusion reads originate but have premature stop codons before the CDS (**Figure 4**).

**Figure 4.**
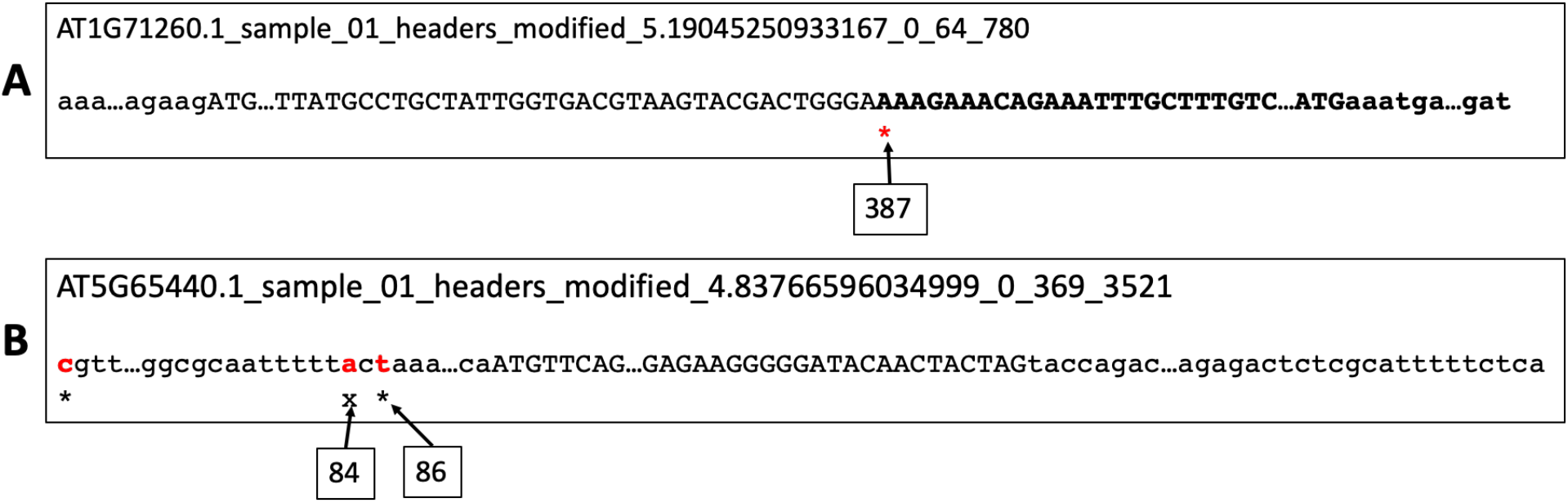
File to assist in primer design. NGPINT generates a text file to design primers for cloning transcripts. Each transcript is represented by three lines. The first line contains the ID of the transcript followed by the sample name where it was detected. This is followed by the fold change, p-adjusted value and start and stop of the coding region. The entire nucleotide sequence is represented in the following line. Both 5’ and 3’ UTR is represented in small case whereas the nucleotides in the coding region are represented by upper case. The final line contains a string from the alphabet {‘*’,’X’, ‘ ‘}. Nucleotides from where fusion reads originate are marked with an asterisk ‘*’ if they are expressed *in-frame*. If fusion reads arise from nucleotides in the 5’ UTR but have a premature stop-codon in the UTR, they are represented by an ‘X’. If no fusion reads originate from a nucleotide, then no symbol is printed beneath it. (**A**) AT1G71260.1 is enriched in-frame and fusion reads are mapped to location 387, which is within the coding region. No fusion reads originate from the 5’ UTR. Portion of the transcript expressed in-frame have been bolded. (**B**) All fusion reads originate from the 5’ UTR regions from locations 1, 84 and 86. Fusion reads starting from position 1 and 86 are in-frame with the coding sequence. Those originating from position 84 will have premature stop codons before the CDS.

## RESULTS

### I. Simulated datasets

To test the robustness of our pipeline, we simulated data from *Arabidopsis thaliana* since most gene models are well-established and complete. We used simulated data because it is impossible to know all true and false positives in an actual experiment. For the simulation, five baits were selected, each with its own unique interaction properties. To represent interacting preys, a set of transcripts were simulated to be differentially enriched. This set of overexpressed transcripts were considered true positives for each bait. Any other transcripts not belonging to this set but reported by NGPINT, were deemed false positives. Often overlapping genes, transcripts of the same gene or genes with similar sequence are incorrectly detected as putative candidates of interaction. Using Y2H-SCORES (Velásquez-Zapata et al., 2020) users can assign priorities to each interacting candidate which is designed to penalize a candidate if there is any evidence of it being a false positive. False negatives comprise those interacting isoforms which were simulated to be differentially enriched but was not reported by NGPINT.

Short reads were simulated using the Polyester package (Frazee et al., 2015). Originally developed to simulate RNA-Seq reads, Polyester generates two samples per replicate – one sample corresponding to the control and the other sample containing higher/lower proportion of reads from differentially expressed transcripts. In the experiment described below, we considered the control sample as the non-selected condition and the other sample as the selected condition. Prior to read generation, each transcript was flanked with the Y2H plasmid sequence. We also added enriched transcripts that were truncated, out of frame, and without vector sequence. A series of fold change values for each transcript (**Supplementary Table 2**), were provided to these scenarios as input to Polyester.

Additionally, we tested parameters that could vary across different sequencing platforms and conditions. Read length, dispersion and error rate were altered to generate reads under different conditions. Illumina error rates were estimated to be 0.24 +/- 0.06% per base (Pfeiffer et al., 2018). Libraries with reads containing base-call errors, ranging from 0.0001 per base to 0.05 per base, were simulated to test robustness. Four different libraries with read lengths 75, 100, 150 and 200 were simulated to see if higher read length could assist in better discovery of interacting candidates. Three replicates for each experiment was simulated. Replicates help in attaining superior estimates of dispersion leading to correct determination of fold change. In this experiment, we tested NGPINT with five different values of dispersion. In addition, we also simulated data at different coverages and varied the minimum trimmed length option to run NGPINT (**Supplementary Table 1**). Finally, NGPINT was executed with both paired- and singleend reads.

To assess the performance of NGPINT, we used an F1 score and Mathew’s correlation coefficient (MCC). F1 score is the harmonic mean of precision and recall. Higher F1 scores signify better recognition of true interactions and fewer false interactions. MCC is a correlation coefficient between the true and predicted binary classes where a higher value indicates better correlation. MCC ranges from −1 to +1 and is suitable for several applications in bioinformatics where unbalanced class sizes exist.

Results from the simulations are summarized in **Table 1**. Transcripts that had a log2 fold change over 1, a p-adjusted value less than 0.01, and *in-frame* reads in at least two replicates were considered as putative candidates. The predicted candidates were compared with the ground truth as defined by our simulated reads to estimate the rate of false discovery. Differentially enriched transcripts simulated in the first mock bait represent an ideal situation where there were no truncated transcripts, no transcripts without vector sequence and all transcripts were expressed *in-frame*. No false positives were detected in this case and 95% of true positives were identified. Partial transcripts were differentially enriched in bait2 (**Supplementary Table 2**) and they were captured appropriately (**Figure 3**). In bait3, bait4 and bait5 transcripts that were out-of-frame were differentially enriched. Several transcripts overlapped with other genes and/or transcripts that were intentionally overexpressed in bait3 and bait5. This sequence similarity resulted in an increase in read count that led DESeq2 to declare those transcripts as differentially abundant. For example, two out of 12 true interactions were missed in bait3, which decreased the recall to 0.83. NGPINT was able to recognize those transcripts and did not declare them as putative candidates, even when some had very high abundance (**Supplementary Table 2**). Analysis of experimental data below (Erffelinck et al., 2018) revealed differential enrichment of several transcripts which had no flanking vector sequence. Transcripts without vector were differentially enriched in bait3, bait4 and bait5. They were recognized as being differentially enriched by NGPINT but were not reported as potential interactors since they are lower priority candidates. Detailed description for each bait and transcript output by NGPINT is illustrated in (**Supplementary Table 3**).

**Table 1.**
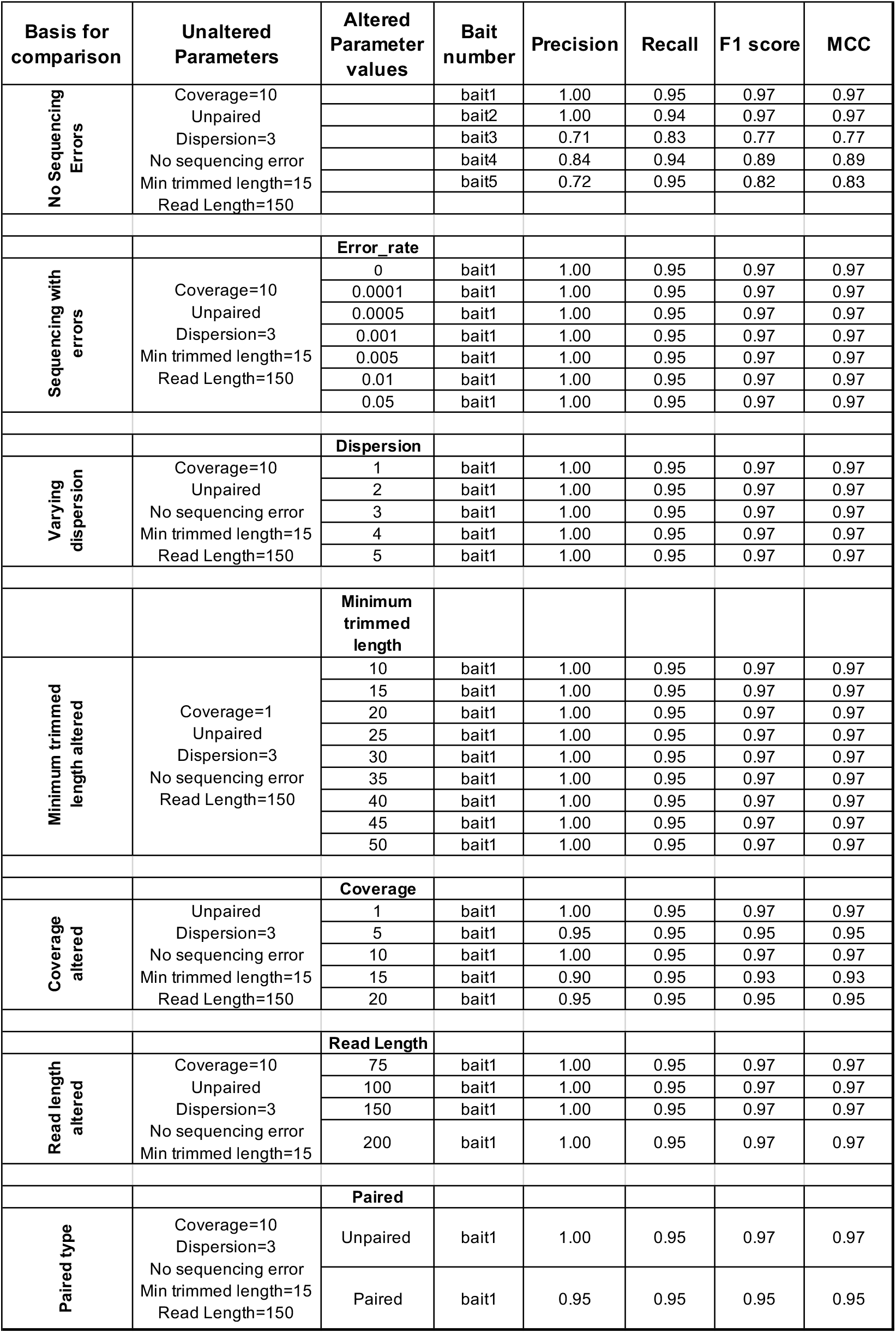
Comparison between ground truth from Arabidopsis thaliana and predicted candidates of protein interaction under different simulated conditions

To summarize, simulations were carried out by varying dispersion, read coverage, read length and error rate. NGPINT was able to detect all interacting candidates in all conditions. For all the baits under different experimental conditions, almost all candidates were recalled successfully. Several genes bear close homology to other genes. Reads from these differentially enriched genes will map to all paralogs and create the illusion of interaction. This impacts precision since it increases false positives. Secondary validation is hence recommended to select the true positives.

#### A. Impact of sequencing errors

Modern sequencing technology seldom produce errors in base calling (Pfeiffer et al., 2018) but are not completely free of inaccuracies. Since the detection of fusion reads is crucial to confirm *in-frame* expression, we simulated datasets with intentional sequencing errors. As illustrated in **Figure 5**, an increase in sequence error rate resulted in a decrease of both fusion-read precision and fusion-read recall. The reduction is expected, since it becomes difficult for NGPINT to predict fusion reads bearing erroneous nucleotides at the junction between the TF and the transcript sequence. But the drop in the detection of fusion reads does not impact the accurate recognition of candidate interactors (**Table 1**) since there are other reads to compensate for those which bear sequencing errors.

**Figure 5.**
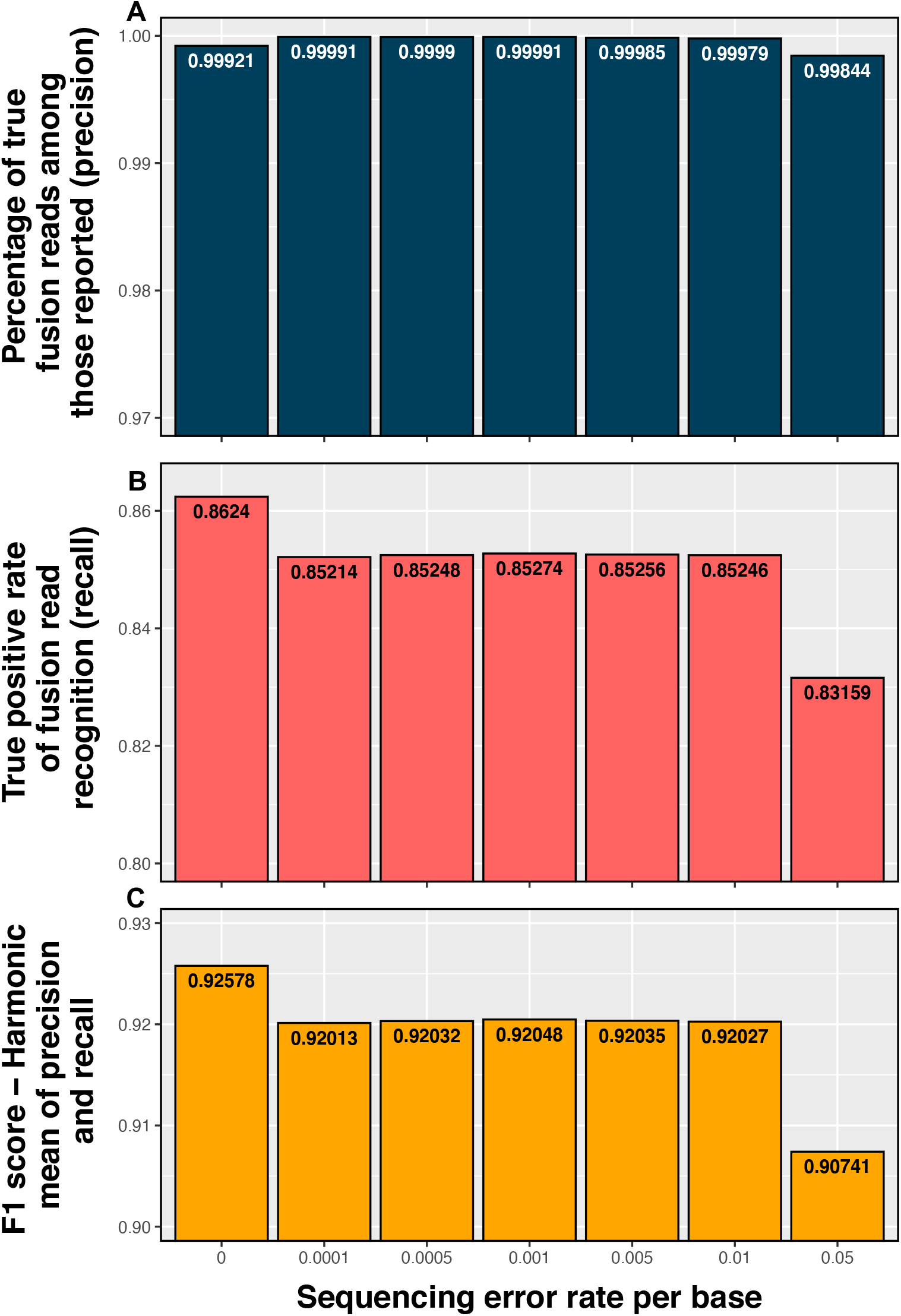
Change in performance of fusion read detection with increasing error rate of sequencing. Very high fraction of fusion reads is correctly recognized when sequencing error is low. The proportion of true fusion reads among all reported fusion reads (precision) remains consistent with increasing error rate. The rate of recognition of true fusion reads (recall) reduces as error rate increases.

#### B. Impact of changing minimum trimmed length

Fusion reads are hybrid sequence fragments that contain both plasmid sequence and organism cDNA. Random shearing of the amplicons prior to sequencing will produce some reads that contain a much lower proportion of vector sequence as compared to organism cDNA. After trimming, such fusion reads can yield very small sequences (<25 bp) of the vector which is not necessarily sufficient to ascertain the actual existence of vector sequence. Hence, a minimum read length is imposed to accurately detect of fusion reads. A similar situation arises when the cDNA fragment is less than the provided threshold. Smaller cDNA fragments will map to multiple transcripts, resulting in an increased read count. Therefore, if a read is detected with a vector sequence or cDNA fragment less than this minimum length (default set to 25), then the read is not classified as a fusion read. We ran NGPINT with different values of minimum trimmed length. As the minimum length increased, fewer fusion reads were being recalled, while the precision remained same **(Figure 6)**. However, this reduction in the number of detected fusion reads did not impact detection of true interacting candidates (**Table 1**).

**Figure 6.**
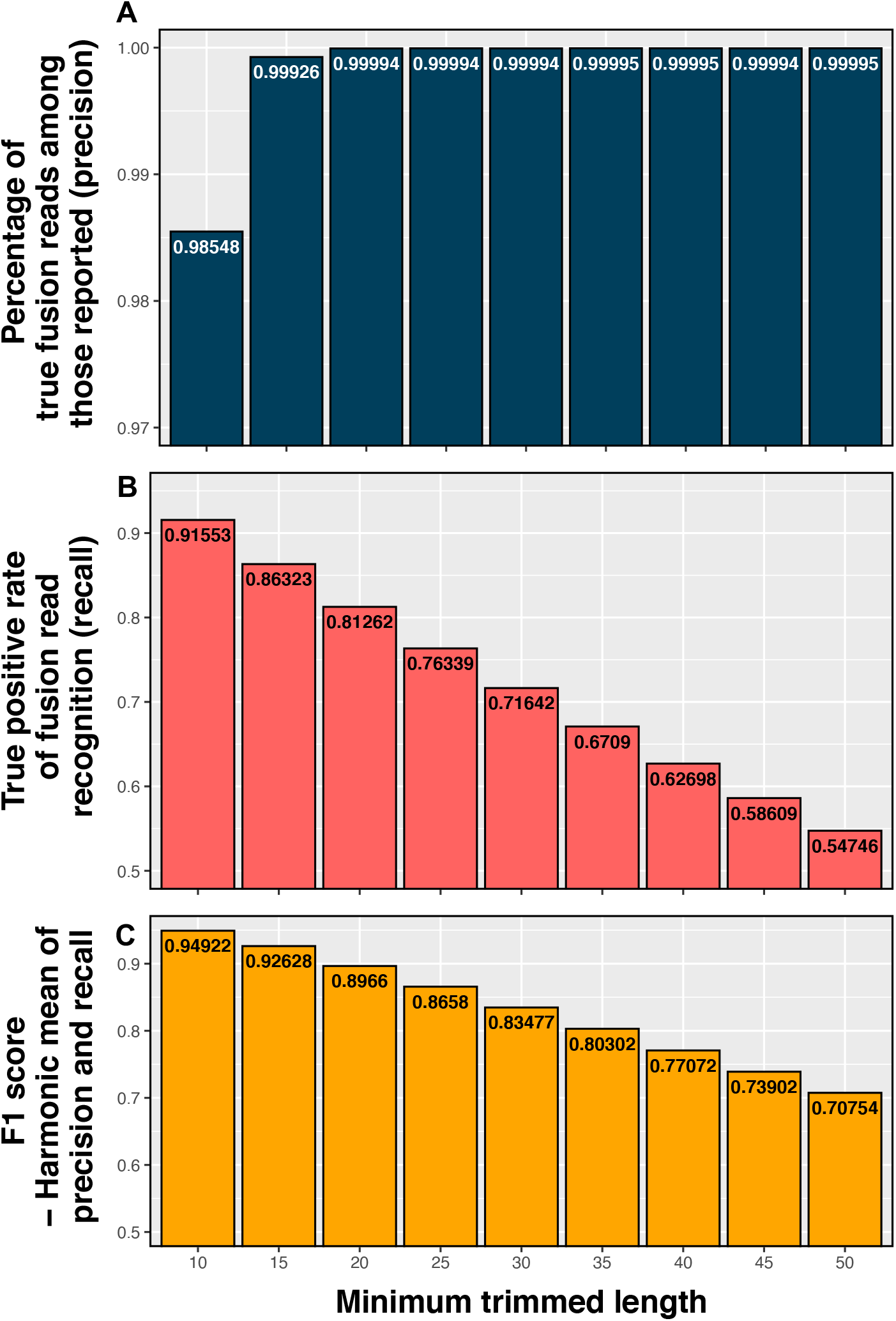
Change in detection of fusion reads with increasing minimum trimmed length. Minimum length of vector portion of soft-clipped reads used as a cutoff to determine fusion reads. Increasing the cutoff leads to recognition of fewer fusion reads but does not impact precision.

#### C. Additional test cases

NGPINT was able to report a very high fraction of true positives (>95%) in all the different categories of simulated data. NGPINT was able to report most of the true positives even when the gene counts were simulated to be highly dispersed (**Table 1**). Library size normalization was implemented which led to the adjustment of the highly dispersed read counts. Hence, DESeq2 was able to detect differentially enriched transcripts. Varying the read length or the coverage did not impact detection of interacting candidates (**Table 1**) showing that the NGPINT performs well with shorter reads and low coverage. The difference noted in precision for different coverages is due to the stochasticity of simulating reads by Polyester. Finally, for both paired and single ended reads, NGPINT was able to report a similar performance – an F1 score more than 0.95.

### II. Testing NGPINT with published datasets

In addition to simulations, we used available experimental data (Erffelinck et al., 2018) to test the performance of NGPINT. Erffelinck and associates (2018) used as bait a previously identified interactor of the jasmonate signalling cascade known as NINJA (AT4G28910) (Pauwels et al., 2010). The Y2H experiment was performed with one replicate and empty vector was used as a negative control. NGPINT was launched with the selected sample from the bait NINJA and the empty vector as background. NGPINT was able to recognize 42 genes which were detected to be highly differentially enriched as compared with the empty vector (Signal-to-Noise ratio (SNR) > 2) and also *in-frame* with the Gal-4 TF (**Supplementary Table 4** [rows 12-53]). Additionally, NGPINT was able to find all 7 previously known NINJA interactors (**Supplementary Table 4** [rows 5-11]) and also detect the 4 newly reported interactors (**Supplementary Table 4** [rows 1-4]) identified by NGPINT. In order to take full advantage of NGPINT and its companion statistical software, Y2H SCORES (Velásquez-Zapata et al., 2020), the experimental design should include multiple replicates. Nevertheless, when NGPINT was executed with the data provided by Erffelinck and associates (2018), the output file successfully returned gene counts, SNR and other relevant information (as outlined in **III. Pipeline outputs**).

The Y2H-NGIS protocol introduces additional unique aspects, for example, expression of overlapping genes or partial transcripts, which require careful consideration. Each of these cases are discussed in detail below.

#### Selecting transcripts for secondary validation

Most eukaryotic genes have multiple transcript isoforms. Often these transcripts share the same sequence, depending on splice junctions. Even when one transcript encodes a protein that interacts with the bait, sequence similarity will prompt the aligner to map reads to all the transcripts. However, it is possible to compare the structures of different transcripts of the same gene, in order to eliminate those transcripts that were not detected. Subtle differences among the transcripts, like presence of unique exons, can be used to select the candidate transcripts. For example, reads originating from the *Arabidopsis thaliana* gene AT5G49540, a Rab5-interacting family protein, map to both transcript isoforms (**Figure 2A**). Both the transcripts share 3 exons but isoform AT5G49540.1 contains an additional 4^th^ exon. Absence of read coverage on the 3rd exon of the AT5G49540.2 isoform and coverage of the 4^th^ exon of AT5G49540.1 is a clear indication that only AT5G49540.1 was detected. Thus, this analysis can prioritize transcript isoforms for secondary validation. NGPINT was also able to detect such cases from the simulated experiments (**Figure 2B**).

#### B. The case of overlapping transcripts

Despite the large size of eukaryotic genomes, genes can be co-located in the same genomic region and overlap. Reads originating from overlapping regions map to both transcripts boosting the read count for transcripts whose protein did not interact with the bait. Read count estimators like Salmon adjusts read counts for transcripts that are only partially covered by reads. One such example is the *A. thaliana* gene AT5G17280, which is present in the 3’ UTR region of a second gene, AT5G17290. The coverage plot indicates that only AT5G17280 is a putative interactor for NINJA. However, read counts for AT5G17290 are artificially increased and also register a high SNR (**Supplementary Figure 2**). Hence, careful analysis of the coverage plots should be performed before proceeding with any further secondary validations.

#### C. The case of partial transcripts

Y2H-NGIS utilizes a cDNA library derived from mRNA isolated from a species and condition of interest. Due to the stochastic nature of cDNA synthesis, this process does not always reverse transcribe full-length cDNAs for all transcripts, resulting in partial cDNAs. However, due to the modular nature of proteins, this can be advantageous in Y2H-NGIS because it allows the user to narrow down specific protein domains that interact with the bait of interest. As shown in (**Figure 3A**), NINJA appears to interact with a protein domain present in the last 40% of the chaperone protein AT5G20890. NGPINT was also able to detect differential abundance of truncated transcripts with simulated data (**Figure 3B**).

## DISCUSSION

Recent developments in sequencing technology have enabled the rapid increase in the availability of high-quality genomes for many non-model species, including plant species of agricultural importance (Van Dijk et al., 2014). Accurate interpretation of proteinprotein interactions underlying signaling networks in these species is key to breeding crops with enhanced stress tolerance. Y2H-NGIS offers a fast, accurate and costeffective approach to discover PPI and map protein interactomes in both model and under-studied organisms (Lewis et al., 2012; Weimann et al., 2013; Pashkova et al., 2016; Trigg et al., 2017; Erffelinck et al., 2018). Although numerous technical advancements have been achieved in Y2H-NGIS protocols, a generalized data analysis pipeline applicable to any biosystem is needed to robustly identify candidate interacting proteins. Here we describe NGPINT - a software package that automates the entire process of analyzing data from Y2H-NGIS protocols. Once executed, NGPINT can run without manual intervention and is configured to run both on Mac and Linux, requiring minimal installations. It is optimized to execute its operations in parallel on the number of permitted CPU cores and can be run on HPC clusters. For example, our analysis using the NINJA dataset (Erffelinck et al., 2018) was completed in less than 1 hour.

Sequence libraries resulting from Y2H-NGIS possess two unique attributes: 1) fusion reads which comprise a hybrid of the prey plasmid and the desired enriched transcripts; these contain junction information to identify transcripts expressed *in-frame*, and 2) high variability of gene counts within replicates.

### A. Fusion reads

NGPINT detects fusion reads by two rounds of alignments of the input prey sequences using STAR (Dobin et al., 2013); fusion reads detected in the first round are trimmed and realigned in the second run, resulting in optimum number of reads aligned to the reference. Accordingly, since cDNA fragments are randomly sheared prior to sequencing, some of these fusion reads contain very short vector sequence or cDNA fragments. This can pose a challenge when using single-end data given that such a small sequence may not be sufficient to align to its reference. Alternatively, paired-end data do not suffer from this issue because mapping of the mate pair can be used to determine the transcript from where the fragment had originated.

### B. Gene Counts

Alignments to the transcripts are generated from the genomic mappings and Salmon (Patro et al., 2017) was used to estimate gene counts. DESeq2 (Love et al., 2014) was used to estimate differential abundance, however, library size normalization (Dillies et al., 2013) was used as opposed to the default median-of-ratios normalization (Anders and Huber, 2010). This is because Y2H-NGIS is expected to exhibit enrichment of all the identified preys in the selected samples. However, this violates the assumption of the median-of-ratios method that requires only a small proportion of the genes to be differentially enriched. In the Y2H-SCORES software workflow, Velásquez-Zapata and associates (2020) showed that using library size normalization over median-of-ratios method has significantly reduced variation of read counts among replicates, thereby improving the power of the statistical tests.

### C. Summary

NGPINT displays exceptional performance when it was executed with data which was simulated with different kinds of errors proving its robustness. NGPINT was able to recall 95% of the true positives when sequencing errors were intentionally introduced in the read data. Difference in read lengths or the choice of paired-end over single-end sequencing did not impact the performance of the pipeline. NGPINT was able to recall most of the true positives even at high dispersion of read counts among the replicates. NGPINT could also recognize transcripts that were (1) differentially abundant but were expressed out-of-frame (2) transcripts with no flanking plasmid sequence and (3) transcripts not having read coverage across its entire length.

One drawback of high-throughput PPI techniques is the presence of auto-activating preys that lead to detection of spurious PPI. One method to detect such cases is to screen many baits and discard those preys that interact with all the baits. If the study is restricted to a few baits, background controls (e.g., empty vector) could be screened to identify and remove the auto-activating preys (Suter et al., 2015). Also, use of the companion software, Y2H-SCORES (Velásquez-Zapata et al., 2020), offers a comprehensive framework to evaluate large-scale PPI screens.

NGPINT offers a user-friendly and cross-platform analysis to detect PPI from NGIS datasets without the need for any specialized computer hardware support. This tool can be further integrated into other complex computational pipelines. Streamlining different bioinformatics tools into a single, easy to use software that addresses unique aspects of Y2H-NGIS data will facilitate rapid PPI mapping in any system under study.

## AVAILABILITY OF CODE, DATA, AND MATERIALS

The python and R scripts code including the software manual file for NGPINT software is provided at GitHub (https://github.com/Wiselab2/NGPINT). Additionally, we implemented a python script to link the Y2H-SCORES functions (Velásquez-Zapata et al., 2020) with the NGPINT pipeline, which will facilitate the integration of both software packages. Using this, it is possible to run the Y2H-SCORES from the NGPINT outputs. Codes and instruction to use Y2H-SCORES can be obtained from (https://github.com/Wiselab2/Y2H-SCORES). Please report any issues regarding the NGPINT software at https://github.com/Wiselab2/NGPINT/issues.

## SUPPLEMENTARY DATA

**Supplementary Figure 1**. Overview of prey insert amplification and sequencing process

**Supplementary Figure 2**. Overlapping transcripts causing increase in read counts.

**Supplementary Table 1**. Description of the arguments of NGPINT

**Supplementary Table 2**. Fold change provided to Polyester for simulation

**Supplementary Table 3**. Detailed description of each interaction reported by NGPINT on the simulated data

**Supplementary Table 4**. Detailed description of each interaction reported by NGPINT on the NINJA data

## ACKNOWLEDGEMENT

The authors thank the Alain Goossens laboratory, Ghent University, Belgium for access to the recent Y2H NINJA-NGIS dataset (Erffelinck et al., 2018).

