## Supplementary figures and images for "NGPINT: A Next-generation protein-protein interaction software"

### Figure S1

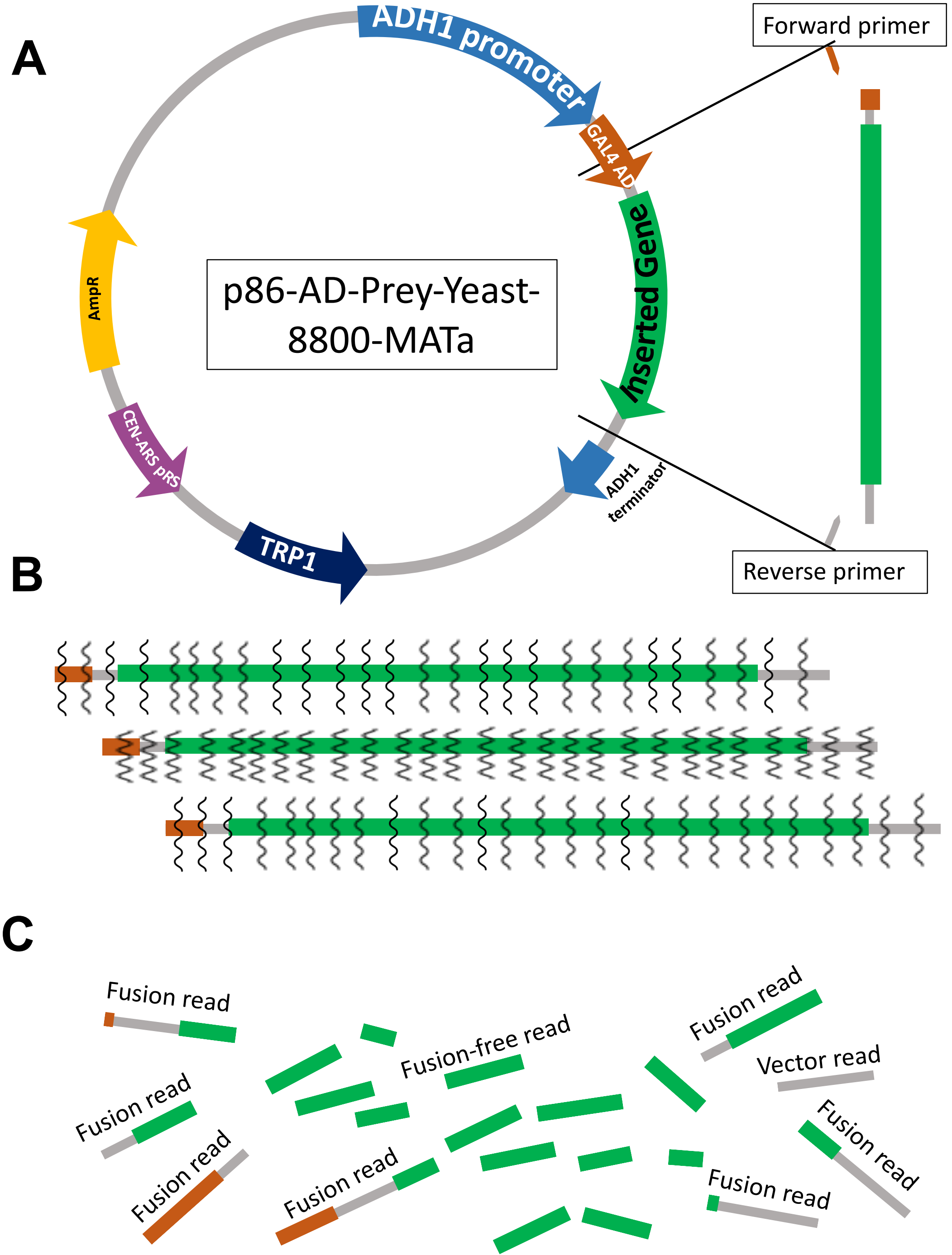

### Figure S2

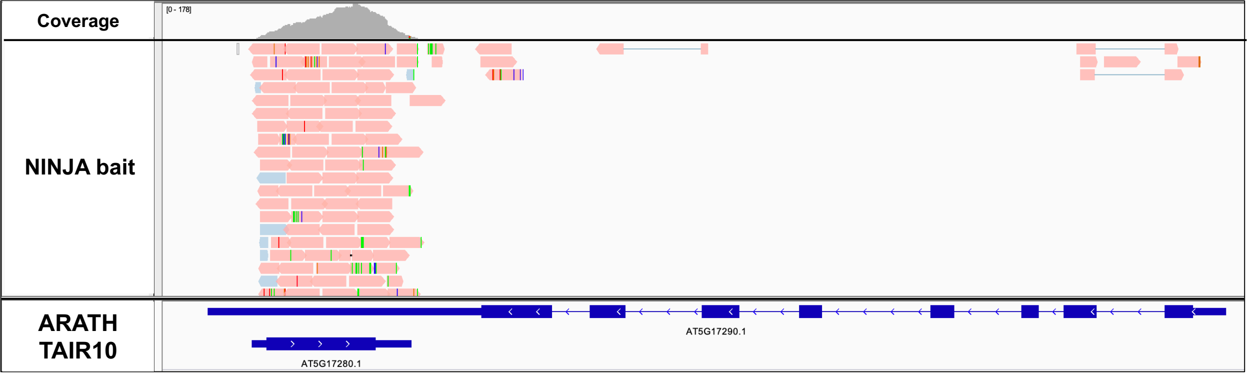
